# Androgen regulation of bowel function in mice and humans

**DOI:** 10.1101/2020.10.15.341081

**Authors:** Daniella Rastelli, Ariel Robinson, Lynley T. Matthews, Kristina Perez, William Dan, Peter Yim, Madison Mixer, Aleksandra Prochera, Rafla Hassan, Kathryn Hall, Sarah Ballou, Judy Nee, Anthony Lembo, Meenakshi Rao

## Abstract

Many digestive disorders have prominent sex differences in incidence, symptomatology, and treatment response that are not well understood. Irritable bowel syndrome (IBS), for example, affects approximately 10% of the population worldwide and tends to have different manifestations in males and females. Androgens are steroid hormones present at much higher levels in post-pubertal males than females and could be involved in these sex differences, but their normal functions in the bowel are largely unknown. Here, we show that gonadal androgens are required for normal gastrointestinal motility *in vivo*. In the healthy mouse gut, we detected androgen receptors in smooth muscle cells and a subset of enteric neurons. Surgical or genetic disruption of androgen signaling in adult mice selectively and reversibly altered colonic motility by affecting neurons rather than smooth muscle. To determine if androgens also influence human bowel function, we measured androgen levels in 208 adults with IBS. Free testosterone levels were lower in patients with IBS compared to healthy controls and inversely correlated with symptom severity. Taken together, these observations establish a role for androgens in the regulation of colonic motility and link altered androgen signaling with a common digestive disorder. These findings advance the fundamental understanding of gut motility, with implications for normal aging and disorders involving the gut-brain axis.

## Main Text

Biological sex affects gene expression at the cell and tissue levels to a surprising degree^1^. These effects could underlie sex differences in the prevalence and clinical presentation of diverse human diseases. Almost all of the disorders of the gut-brain axis, from inflammatory bowel disease to functional gastrointestinal (GI) disorders, and even neurological disorders with GI comorbidities, such as Parkinson’s disease, exhibit prominent sex differences^2–4^. Females are disproportionately affected by some of these disorders in Western countries leading to previous exploration of estrogens as a risk factor in digestive disease. This work has shown that estrogens can modulate intestinal epithelial barrier function^5^, smooth muscle contractility ^6^, and enteric neurogenesis after injury^7^. Androgens, such as testosterone and 5α-dihydrotestosterone (DHT), found at much higher levels in post-pubertal males than females, have received less attention and their role in GI homeostasis remains poorly understood.

Androgens signal through androgen receptor (AR), a ligand-dependent transcription factor, to activate changes in gene expression. In addition to this canonical pathway, androgens also have non-genomic effects that are faster and independent of gene transcription^8^. Testosterone and DHT are the primary ligands for AR, but DHT is 10-fold more potent, and unlike testosterone, cannot be aromatized into estradiol. Previous work demonstrated that radiolabeled DHT binds cell nuclei throughout the GI tract of non-human primates, predominantly in the muscularis externa^9^, suggesting that intestinal smooth muscle is androgen-responsive, similar to vascular smooth muscle. Consistent with this, smooth muscle strips from the small and large intestines of mice exhibit increased contractility when pre-incubated with androgens *in vitro*^10,11^. It is unknown, however, to what extent androgens regulate bowel functions *in vivo*.

To determine the normal functions of androgens in digestive health and gut motility, we performed bilateral orchiectomy (surgical castration, hereafter referred to as ORCH) in wildtype male mice, shortly after puberty. Four weeks later, we measured GI transit time in ORCH mice and littermate controls that underwent sham operations (same surgical procedure without testes removal, hereafter SHAM). Seminal vesicle weight, a validated measure of androgen signaling activity^12^, was markedly lower in ORCH mice and confirmed the efficiency of androgen depletion (Extended Fig. 1). We found that ORCH and SHAM mice were indistinguishable in their cages with no difference in body weight (Extended Fig. 1), but GI transit time was 40% slower in ORCH mice (Fig. 1a). This dysmotility persisted with no change in magnitude over the next 16 weeks (Fig. 1b). ORCH mice also had 25% larger fecal pellets with less water content (Fig. 1c,d), additional features of chronic constipation. These observations suggest that gonadal function is necessary in adult male mice for normal GI motility and that its loss causes constipation.

**Figure 1.**
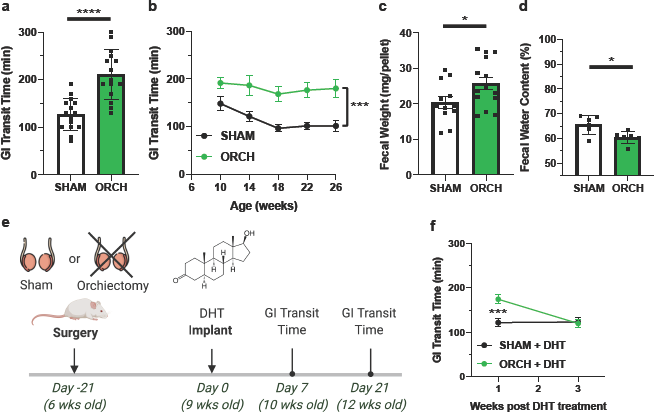
Gonadal androgens are required for normal gastrointestinal transit in vivo. **a.**Total gastrointestinal (GI) transit time was measured in 10-week-old wildtype (FVB/NJ strain) male mice four weeks after sham surgery (SHAM; n = 15) or bilateral orchiectomy (ORCH; n = 15). ORCH mice had markedly delayed GI transit compared to SHAM mice (P < 0.0001). **b.**Total GI transit time was measured every 4 weeks after surgery and remained consistently slower in ORCH mice compared to SHAM controls (n = 6/group, P = 0.001, 2-way repeated measures ANOVA). **c.**Average fecal pellet mass was greater in ORCH mice than SHAM controls (n =12-14/group; P = 0.0433). **d.**Fecal water content was lower in ORCH mice than SHAM controls (n = 6/group; P = 0.0198). **e.**Schematic of experimental design for androgen rescue experiment. Bilateral orchiectomy or sham surgeries were performed on 6-week-old FVB/NJ mice. Subcutaneous pellets that continuously released 5α-dihydrotestosterone (DHT), a non-aromatizable androgen, were implanted into mice from both groups three weeks later. Total GI transit time was then measured at 10- and 12-weeks of age. **f.**Three weeks of DHT supplementation resulted in the normalization of GI transit time in ORCH mice to that of SHAM mice (P = 0.0029, 2-way repeated measures ANOVA). * Represents P < 0.05, * * P < 0.05, * * * P< 0.005, and * * * * P<0.001. Unpaired t-tests used to compare pairs of group means in **a, c** and **d**.

The testes are the predominant source of circulating androgens in males but also elaborate other bioactive molecules. To ascertain to what extent androgen deficiency was responsible for slowed GI transit in ORCH mice, we restored androgens three weeks after surgery by implanting subcutaneous pellets that continuously eluted 0.125 mg/day of DHT (Fig. 1e). Supplemental DHT did not affect GI transit time in SHAM mice, but completely normalized the GI transit deficit in ORCH mice within three weeks, suggesting that androgens were sufficient to rescue the dysmotility caused by orchiectomy (Fig. 1f).

GI motility represents a complex repertoire of motor behaviors that vary along the digestive tract. To determine which motor behaviors are androgen-dependent, we analyzed gastric, small intestinal, and colonic motility in ORCH mice to localize the deficit. We found that gastric and small bowel transit were unchanged in ORCH mice (Fig. 2a, b), suggesting that dysmotility localized to the colon. To assess the gut-intrinsic effects of androgen deficiency, we studied colonic motility *ex vivo* in full-length colonic segments acutely isolated from SHAM and ORCH mice, four or more weeks after surgery. The neural circuits required for colonic peristalsis are part of the enteric nervous system (ENS) intrinsic to the GI tract^13^. As such, rhythmic colonic migrating motor contractions (CMMCs) that propel luminal contents in the expected oral to anal direction can be observed *ex vivo*^14,15^. We imaged colonic motility in SHAM and ORCH mice and found that colons from ORCH mice exhibited highly disorganized contraction patterns (see videos in Extended Fig. 2) with fewer effective CMMCs that propagated at least 50% of the length of the bowel (Fig. 2c-e; Extended Fig. 3). These observations indicate that loss of gonadal function in male mice selectively disrupts colonic motility.

**Figure 2.**
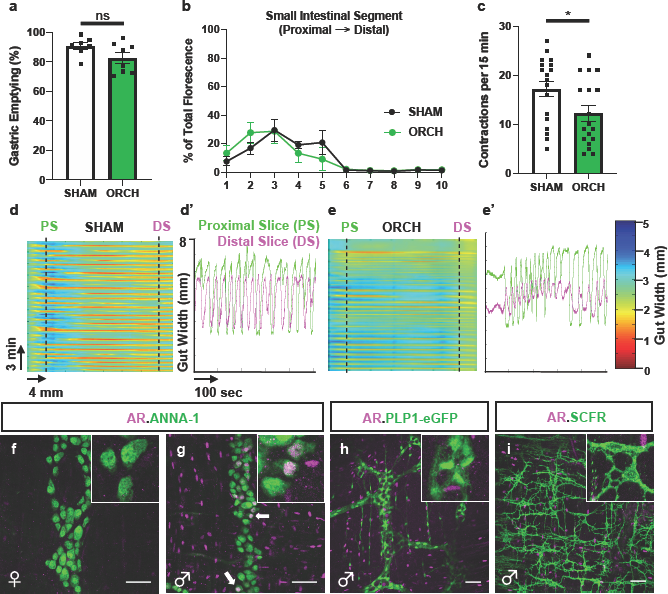
Gonadal androgen deficiency selectively disrupts colonic motility. **a.**Gastric emptying of a non-absorbable dye was no different between SHAM and ORCH mice (n = 7-8/group; P = 0.0967). **b.**Small intestinal transit, measured as the distance traveled by a fluorescent dye along the small intestine 15 minutes after gavage, was no different between SHAM and ORCH mice. X-axis represents intestinal segment number (proximal à distal). **c.**Ex vivo imaging showed that colons from ORCH mice had fewer effective, colonic migrating motor contractions (defined as those starting proximally and propagating > 50% of the length of the bowel), than those from SHAM mice (* P = 0.0150). **d,e.**Representative spatiotemporal maps illustrating contractile activity of colons from SHAM (**d**) and ORCH (**e**) mice ex vivo. Color reflects degree of gut contraction (measured as gut width). Vertical slice analysis at proximal (PS) and distal segments (DS) of the SHAM mouse colon (**’**) shows regular periodic contractions that start proximally and propagate distally (each green drop in gut width is followed by a pink drop). Similar analysis of ORCH mouse colon (**e’**) shows more disorganized contractions with less efficient propagation. See Extended Data Figure 2 for videos. **f,g.**Androgen receptor (AR) immunoreactivity is undetectable in the muscularis externa of the colon from an adult female mouse (**f**), but abundant in that of an adult male (**g**). The majority of AR-expressing cells are in the smooth muscle but a subset colocalizes with the pan-neuronal marker (ANNA-1; arrows) in the myenteric plexus. **h.**Enteric glia labeled with green fluorescent protein (GFP) in the colon of an adult male PLP1-eGFP mouse do not express AR. **i.**Interstitial cells of Cajal labeled by SCFR immunoreactivity do not express AR in the adult male mouse colon. Scale bars in **f–i** = 50μm. Insets in **f-i** show representative cells from associated images at higher magnification.

**Figure 3.**
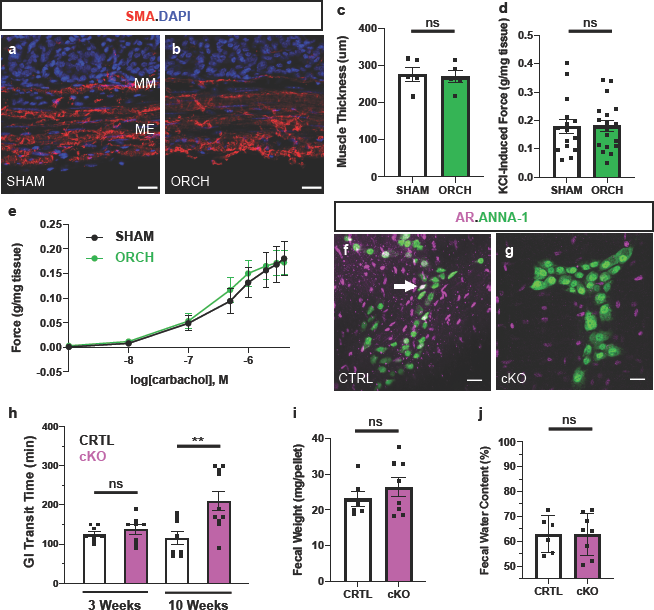
Androgen effects on gut motility are mediated by signaling in neurons, rather than smooth muscle. **a,b.**Cross-sections of colons from SHAM and ORCH mice immunostained for α-smooth muscle actin (SMA) show that the muscularis externa (ME) and muscularis mucosa (MM) are grossly normal in ORCH mice. Cell nuclei marked with DAPI. Scale bars = 20μm. Tissues for analyses in **a-e** were isolated from mice 5-7 weeks after surgery. **c.**There is no difference in colonic smooth muscle thickness between SHAM and ORCH mice (n = 5 mice/group; P=0.7824). **d.**Maximum contractile force exerted by colonic segments from SHAM (n = 15 segments from 5 mice) and ORCH mice (n = 19 segments from 6 mice) is no different (P = 0.9503). Force measurements in **d** and **e** were normalized to tissue mass. **e.**The dose-dependent contractile force generated by colonic segments upon exposure to the acetylcholine receptor agonist carbachol, is no different between segments isolated from SHAM and ORCH mice (n = 15-19, as in **d**). **f,g.**Androgen receptor (AR) and pan-neuronal marker (ANNA-1) immunoreactivities show that AR expression is evident in both myenteric neurons and surrounding smooth muscle in colons from 10 week-old AR^flx/Y^ control mice (CTRL), but undetectable in enteric neurons of AR^Wnt1KO^ mice (cKO). Arrow marks an AR^+^ neuron in the CTRL colon. Scale bars = 25 μm **h.**GI transit time was no different in pre-pubertal (3-week-old) CTRL and cKO mice. When re-assessed at a post-pubertal age (10 weeks), GI transit time in cKO mice was prolonged (n= 8-10 mice/group; P = 0.0027). **i,j.**Fecal pellet mass (**i**) and water content (**j**) were no different between CRTL and cKO mice at 10 weeks of age.

Normal colonic motility arises from the coordinated actions of many cell types, including smooth muscle cells, interstitial cells of Cajal (ICCs), enteric glia, and enteric neurons, all found within the muscularis externa^16–18^. To determine which of these cell types might mediate the effects of androgens on colonic motility, we examined the expression of AR by whole-mount immunohistochemical staining of colonic segments from wildtype, post-pubertal male and female mice. AR was undetectable in the muscularis externa of female mice (Fig. 2f) but was found in a large population of cells in males (Fig. 2g). Nuclear AR-immunoreactivity was evident in many smooth muscle cells, consistent with previous reports; unexpectedly, it was also present in the myenteric plexus of the ENS, where it co-localized with the pan-neuronal marker ANNA-1 (Fig. 2g). Remarkably, 17 ± 1.7% of neurons within the myenteric plexus, but none in the submucosal plexus, expressed AR (mean ± SEM; *n* = 4 mice). AR was not detected in enteric glia or ICCs (Fig. 2h, i). These data suggest that in addition to intestinal smooth muscle, a subset of enteric neurons in the adult male colon can respond to androgens.

The colonic dysmotility caused by loss of gonadal function and androgen deficiency could be due to a requirement for androgen signaling in intestinal smooth muscle, enteric neurons, or both. Smooth muscle bulk, measured by immunohistochemical staining for alpha-smooth muscle actin, was no different between SHAM and ORCH mice (Fig. 3a-c). To test smooth muscle function, we measured the contractility of acutely isolated colonic rings in the presence of tetrodotoxin, which blocks voltage-gated sodium channels and thus most fast synaptic transmission in the ENS^19^. Both maximal contractile force of the smooth muscle, induced by KCl, and dose-dependent contractile force in response to the acetylcholine receptor agonist carbachol, were no different between SHAM and ORCH mice (Fig. 3d, e). These findings show that loss of male gonadal function did not diminish intestinal smooth muscle bulk or contractility and suggest that colonic dysmotility in ORCH mice is instead due to a requirement for androgen signaling in enteric neurons.

The effects of androgens on the nervous system have been most studied in the brain, where they influence its circuits in two distinct phases. During development, a transient surge of androgens produced by the testes establishes sex-specific circuits in certain brain regions, such as the hypothalamus^20^. The second phase occurs upon puberty, when androgens again surge in males and begin to modulate the activity of these and other brain regions important for sex-specific behaviors, including mating^21^. Consistent with this, circulating testosterone levels in male mice exhibit a biphasic pattern over time, with an initial peak within 24 hours of birth and a second much larger peak at five weeks of age, preceding breeding capability by 4-5 days^22^. We found that AR expression was undetectable in the colons of pre-pubertal male mice (Extended Fig. 4), but robust in enteric neurons and surrounding smooth muscle by 7 weeks of age (Fig. 2g), suggesting that the ENS becomes androgen-responsive during puberty. Bilateral orchiectomy after puberty had no effect on neuronal density in the myenteric plexus (Extended Fig. 4d), implying that androgen deficiency affects enteric neuronal function rather than their genesis or survival.

**Figure 4.**
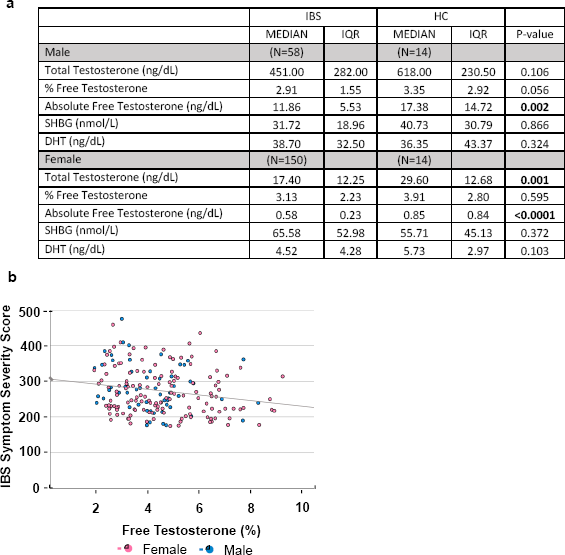
Diminished testosterone levels correlate with the diagnosis and severity of irritable bowel syndrome. **a.**Median values and inter-quartile ranges (IQR) for total testosterone, percent (%) free testosterone, sex hormone-binding globulin (SHBG), and dihydrotestosterone (DHT) measured in the sera of a cohort of post-pubertal adults with irritable bowel syndrome (IBS) and healthy controls (HC). Absolute free testosterone was calculated by multiplying total testosterone by the percent that was free. N = number of subjects. P-values are for t-tests comparing means (% free testosterone) or means of log-transformed values (all others) between IBS and HC groups of each sex. **b.**Scatter plot illustrating the relationship between IBS symptom severity score (IBS-SSS) and percent free testosterone in the sera of patients with IBS. Higher score indicates more severe symptoms. Percent free testosterone was no different between males and females (t-test, P = 0.180), therefore these groups were combined. Partial correlation analyses controlling for age and sex revealed a negative association between percent free testosterone and IBS symptom severity (r = −0.19, P = 0.009).

To determine the extent to which enteric neuronal AR signaling is necessary for normal motility, we generated conditional knockout mice lacking AR in enteric neurons but not in the surrounding smooth muscle. Given the absence of Cre lines specific to enteric neurons, we used the Wnt1^Cre2^ transgenic line, which has been widely utilized to manipulate genes in neural crest derivatives, including the ENS^23,24^. We first characterized Wnt1^Cre2^ activity in the adult mouse colon using Ai9 Cre reporter mice and found that Cre activity was detectable throughout the ENS, including the majority of enteric neurons, but never in the smooth muscle (Extended Fig. 5). Wnt1^Cre2^::Ar^flx/Y^ mice (hereafter AR^Wnt1KO^) were then generated and confirmed to lack AR expression in the myenteric plexus, but not neighboring smooth muscle cells (Fig. 3f, g). AR^Wnt1KO^ mice were born at expected Mendelian frequencies, appeared healthy, had similar body weights to Cre-negative, Ar^flx/Y^ littermate controls (Extended Fig. 6), and were fertile. We measured GI transit time in AR^Wnt1KO^ male mice at three and ten weeks of age to assess the consequences of AR signaling to the ENS pre- and post-puberty. We found that while three-week-old AR^Wnt1KO^ mice had the same average GI transit time as their littermate controls, ten-week-old AR^Wnt1KO^ mice had delayed GI transit, similar to ORCH mice (Fig. 3h). Fecal pellet size and water content, in contrast, were no different in AR^Wnt1KO^ mice (Fig. 3i,j), suggesting that neuronal AR depletion specifically disrupts GI motility. In total, these results show that the ENS becomes androgen-responsive during male puberty and that androgen signaling through AR in the peripheral nervous system is necessary for normal gut motility.

IBS is a common and debilitating functional GI disorder of unclear etiology that is characterized by abdominal pain and altered stool patterns (diarrhea, constipation, or both). Two studies in which testosterone levels were measured by immunoassay in small groups of males with IBS found associations between testosterone levels and GI symptoms^25,26^, suggesting that altered androgen signaling might play a role in IBS. To test this possibility more rigorously, we used a highly-sensitive mass spectrometry assay to measure serum total testosterone, percent free testosterone, and DHT levels in 208 patients with IBS at the baseline visit of a randomized controlled trial designed to test a therapeutic intervention (Clinical Trials.gov NCT02802241)^27^. Absolute free testosterone, the form bioavailable for signaling, was determined by calculating the proportion of total testosterone that was measured as free. We found that both males and females with IBS had lower levels of circulating free testosterone than healthy controls, with no difference in DHT levels (Fig. 4a; Extended Figure 8), suggesting that IBS is associated with selective androgen deficits. To determine whether androgen levels correlated with symptoms, we examined the association between free testosterone and IBS symptom severity score (IBS-SSS), a commonly used composite measure of IBS severity, using partial correlation analyses that controlled for patient age and sex. We focused on percent free testosterone because this is similar in both sexes, while absolute levels can differ by 5-10 fold. IBS-SSS inversely correlated with percent free testosterone in both sexes (Fig. 4b), suggesting that having a lower proportion of bioavailable testosterone is associated with more severe symptoms in patients with IBS.

Androgen levels are markedly different in males and females after puberty, and can be further regulated at the local tissue level, offering a robust substrate for sex differences in health and disease. Here we show that abolishing gonadal androgens in otherwise healthy adult mice causes a profound deficit in gut motility that is fully rescued by restoring a single androgen hormone. Unexpectedly, we found that a population of enteric neurons becomes androgen-responsive upon puberty and that androgen signaling through AR in these and other peripheral neurons is essential for normal gut motility in adult mice. Previous studies of androgen signaling to the peripheral nervous system have largely focused on sex-specific motor functions, such as the penile erectile response^28^. Our findings suggest that androgens have much broader roles in modulating peripheral neuronal activity that merit exploration.

Circulating androgens are eliminated from the body by glucuronidation and excretion into the urine and bile. Bile is secreted into the gut lumen where specific microbes can deconjugate glucuronides, generating high local levels of steroid hormones, especially in the colon^29,30^. This phenomenon could constitute a rich source of ligands for AR-expressing neurons that innervate the gut and explain why colonic motility was particularly vulnerable to the effects of androgen deficiency in our study. Modulating peripheral nervous system function by recycling androgens into bioactive forms could be a mechanism for microbes to modulate host physiology.

After establishing proof of concept of androgen function in gut motility in mice, we investigated androgen levels in humans with IBS, a common GI disorder associated with altered gut motility. Because we leveraged a patient cohort assembled to address a therapeutic question, our study lacked a large number of age-matched, healthy controls and exclusion criteria related to hormone-based therapies. Despite these limitations, we found that altered androgen levels were associated with the diagnosis and severity of IBS in both males and females. Our preclinical studies focused on male mice because we did not detect AR expression in the female colon; however, the IBS data suggest that androgen signaling is likely to be involved in gut functions in both sexes. This could reflect species differences or, more likely, the limits of immunohistochemistry to detect low levels of AR. Consistent with this, a previous study using a genetic reporter suggested that *Ar* is transcriptionally active in the female mouse intestine^31^.

Overall, our findings establish androgen-AR signaling as a critical pathway in the regulation of gut motility and suggest that it should be considered in the pathophysiology of disorders affecting the gut-brain axis. Androgen levels decline with age in men and up to 20% of males over the age of 60 have low levels of bioavailable testosterone^32^. Thus, diminished androgen signaling in the peripheral nervous system is likely to play a role not just in digestive disease, but also in the consequences of normal aging on gut functions.

## Methods

### Mouse lines

Mice were housed in a specific pathogen-free facility with a 12-hour dark cycle and handled per protocols approved by the Animal Care and Use Committees of Boston Children’s Hospital and Columbia University Medical Center. PLP1^eGFP^ mice (JAX 033357; Jackson Laboratory, Bar Harbor, ME) were bred on a FVB/NJ (JAX 001800) background. All other mice were maintained on a C57Bl/6j background. Wnt1^Cre2^ hemizygous males (JAX 022137) were bred with female Rosa26^Ai9^/Ai9 mice (JAX 007676) to generate Wnt1^Cre2^::Rosa26^Ai9/+^ mice to test Cre activity. AR^flx/flx^ (EMMA 02579; Infrafrontier, Munich, Germany) female mice were bred with Wnt1^Cre2^ hemizygous males to generate Wnt1^Cre2^::AR^flx/Y^ mice and AR^flx/Y^ littermate controls.

### Orchiectomy and Sham Surgical Procedures

Six-week-old male FVB/NJ mice were anesthetized with 1-3% inhalable isoflurane (Patterson veterinary NDC 14043-704-05), administered ophthalmic eye ointment (Dechra NDC 17033-211-38), 5-10 mg/kg subcutaneous meloxicam (Putney NDC 26637-621-01), and subcutaneous bupivacaine at the incision site (MWI Veterinary Supply, Boise, ID; 0.25%, 2-3 mg/kg diluted with 50% saline for final 0.125% solution). Fur was removed from the scrotum, and the surgical area was sterilized with chlorhexidine and 70% ethanol. Using sterile technique, a single incision was made on the midline of the scrotum. Blunt forceps were used to grasp one testicular fat pad, and a ligature of absorbable suture material was placed around the epididymis. Using sharp scissors, the testicle was removed, and the remaining tissue returned to the scrotal sac. Using the same incision, the procedure was repeated to excise the second testicle. After ensuring adequate bleeding control, the skin was closed with simple interrupted non-absorbable sutures. For sham operation, the same scrotal incision was made, the testicles were visualized, and then testes were returned to the scrotal sac. All animals received meloxicam 24 and 48 hours after the first injections to minimize postoperative pain.

### DHT Subcutaneous Implant Procedure

Mice were anesthetized and provided with peri-procedural bupivacaine and meloxicam analgesia, as described above. Pellets containing 5-alpha-dihydrotestosterone (DHT) were implanted subcutaneously under sterile conditions (Innovative Research of America). The pellets contained 7.5 mg DHT/pellet and were designed to continuously release 0.125 mg per day of DHT over 60 days. No post-procedure analgesia was administered, as advised by veterinary consultation.

### Immunohistochemistry, microscopy, and cell quantification

Gut tissues were isolated and immunostaining performed as previously described (Rao, 2015). Primary antibodies used were: Human anti-ANNA-1 1:40,000 (Gift from V. Lennon), Rabbit anti-AR 1:250 (Santa Cruz sc-816), Chicken anti-GFP 1:1000 (AVES GFP-1020), Goat anti-SCFR 1:500 (R&D Systems AF1356), Rabbit anti-SMA 1:200 (Abcam ab-5694), Rabbit anti-nNOS 1:1500 (ImmunoStar 24287), Rabbit anti-PGP9.5 1:1000 (Cedarlane CL95101). Anti-goat and anti-donkey secondary antibodies used were conjugates of AlexaFluor 594, 568, or 488 (Invitrogen). Nuclei were counterstained with DAPI in Vectashield mounting medium (Vector Labs H-1200, Burlingame, CA). To quantify AR^+^ neurons, 1.5cm segments of the colon were immunostained as whole mounts for ANNA-1 and AR. Single planar images were captured at the level of the myenteric plexus (6 fields per segment, identified based on ANNA-1 signal alone). Images were coded, randomized, and quantified using ImageJ by investigators blinded to the experimental condition. To quantify smooth muscle thickness in SHAM and ORCH mice, colons were acutely isolated, gently flushed to remove fecal pellets, and then fixed with 4% paraformaldehyde in phosphate buffered saline while cannulated on bamboo skewers, to ensure equivalent luminal distention across all samples. Cryosections of colonic tissue (14μm) were immunostained for SMA, mounted in Vectashield with DAPI, and imaged on an Olympus BX41 epifluorescent microscope. Twelve images were captured per animal and muscularis externa thickness was measured using SMA to mark boundaries. All images of immunohistochemical staining in the manuscript are representative of observations made in a minimum of 3 mice per condition.

### Gastrointestinal Motility and Stool Composition Analyses

Total gastrointestinal transit time, gastric emptying and small intestinal transit were measured as previously described (Rao et al., 2017). For *ex vivo* imaging of colonic motility, colonic contractile activity was recorded and analyzed using Scribble and Matlab scripts, as previously described (Rao et al., 2017; Swaminathan et al, 2016). Briefly, this analysis generates spatiotemporal heat maps in which color represents gut width, the X-axis represents the proximal to distal length of the colon, and the Y-axis represents time. CMMCs were defined as contractions that originated in the proximal colon and successfully propagated at least 50% of the length of the tissue. All parameters were measured from three 15-minute video recordings obtained from each of 6 mice per group, and all individual data points are shown in graphs. The organ bath preparation accommodates 2 colons at a time, and SHAM and ORCH colons were imaged in every session in parallel. To determine the rate and time required for relaxation of individual CMMCs, vertical slice analysis was performed at two positions along the x-axis of the heat map: one at 1/3^rd^ of the length of the colon (proximal slice) and one at 2/3rds of the length (distal slice). This generates line graphs at these two points showing gut width over time. The green line shown in Figure 2d’ and 2e’ represents the vertical slice from the proximal positions and the pink line represents the distal position. From these graphs, the rate of relaxation was calculated from determining the change in gut width over the change in time as averaged from two contractions per video. For stool composition measurements, mice were transferred to clean cages without bedding at 9am and stool output was continuously collected for 1 hour. Animals had free access to both food and water during this time. Total mass of stool collected was “wet” weight per mouse. Fecal pellets were then dried at 45°C for a minimum of 48 hours until a stable mass was reached, and then a “dry” weight was recorded. Percent stool water content = [(“wet”-”dry”)/”wet”]*100). Average fecal pellet mass = “wet”/ # of pellets.

### Smooth Muscle Contractility

Mice were euthanized with 40 mg/kg sodium pentobarbital. The colons were removed and flushed with modified Krebs buffer (mM: NaCl 137, KCl 2.9, CaCl2 1.8, MgCl2 2.1, NaH2PO4 0.4, NaHCO3 11.9, D-Glucose 5.6, pH 7.4). Connective tissue was removed, and 3-4 mm lengths of transverse annular segments of the distal colon were excised ≥1 cm from the anus and massed. Each colonic ring segment was mounted in a chamber of a DMT 620M Multi-Wire Myograph (DMT, Ann Arbor, MI). The Krebs buffer was changed every 15 min (37°C, bubbled at 95% O2/ 5% CO2), and each tissue was equilibrated at 1 gram (g) resting tension for 1 hour. To demonstrate tissue viability, colonic smooth muscle rings were isometrically contracted with three cycles of increasing log concentrations of acetylcholine (ACh) (10nM-1mM) (Sigma Aldrich, St. Louis, MO). Krebs buffer was exchanged, and tension was readjusted to 1g between each ACh cycle to re-establish the baseline tension. After the final ACh cycle, and following three buffer exchanges, all tissues were pretreated with 1 uM tetrodotoxin (EMD Millipore, Jaffrey, NH) to mitigate the pro-relaxant effects of the myenteric nerve plexuses and maximize myogenic contractility. After 20 min pretreatment, the colonic rings were contracted with carbachol 1nM-4uM (Sigma Aldrich, St. Louis, MO) to define the EC50 carbachol of each colonic ring. The buffer was exchanged, and resting tension returned to 1g. Tetrodotoxin pretreatment was re-administered before maximal contraction was induced with 200 mM KCl.

### Human IBS Study and Statistical Analysis

Androgen and sex hormone binding globulin (SHBG) levels were measured in serum collected at the baseline visit of a randomized controlled trial designed to evaluate open label placebo as a therapeutic modality in patients with IBS (Clinical Trials.gov NCT02802241). All subjects in this IRB-approved trial were recruited from a single academic medical center in the United States of America. Hormone measurements were obtained using liquid chromatography tandem mass spectrometry (LC-MS/MS) assays and SHBG was measured using a chemiluminescence immunoassay. Assays were performed by the Brigham Research Assay Core (BRAC; Brigham and Women’s Hospital, Boston, MA), which is certified by the Center for Disease Control’s Hormone Assay Standardization Program (HoST). Measurements were made for all 209 individuals with IBS in the study for whom serum samples were available from the baseline visit and 28 healthy controls (HC). Of these, one individual with IBS was excluded due to an indeterminate measurement. Final analyses were based on the remaining 236 subjects (208 IBS, 28 HC) using SPSS (IBM SPSS Statistics version 27, IBM Corp., Armonk, NY, U.S.A.). Extended Figure 7 reports descriptive data of study subjects, including age and distribution of IBS-subtypes, as means and standard deviations (SD). This study was not powered to detect subtype-specific differences in hormone levels. Median values and interquartile ranges (IQR) of all hormone levels are reported in Figure 4a, and also illustrated as box plots in Extended Figure 8. Because the distributions of the hormone data were skewed, statistical analysis was conducted after transforming this data using a logarithmic scale (with natural log values). This transformation was applied to all measurements except for percent free testosterone because this measure was already normally distributed. T-tests were used to evaluate the differences in mean log-transformed levels between IBS and HC subjects, separately for males and females.

The Irritable Bowel Syndrome-Symptom Severity Scale (IBS-SSS) is a validated 5-question survey used to generate a composite score based on: severity of abdominal pain, number of days with abdominal pain over the preceding 10 days, presence and severity of abdominal distension, satisfaction with bowel habits, and IBS-related quality of life (Francis et al., 1997). Each of these 5 components is scored on a scale of 1-100 with a maximum composite score of 500. Partial correlation analysis was used to assess the association between IBS-SSS and percent free testosterone. Student’s t-test revealed no difference in mean percent free testosterone between male and female IBS patients, so the groups were combined for correlation analysis to increase power, controlling for age and sex (results shown in Figure 4b).

### Statistical Analyses of Non-Human Data

For comparisons between pairs of means, unpaired Student's t-tests were used. All graphs display mean ± standard error of the mean (SEM) for each condition with each individual data point shown. “*n*” refers to number of mice per condition, except where specifically noted. A *P* value < 0.05 was considered significant and denoted with *. *P* values < 0.005 are noted with * * and < 0.0005 with * * *. For comparisons of means from 3 of more groups, ANOVA was used. For repeated measures over time compared between two groups, 2-way ANOVA and post-hoc Tukey tests were performed. For muscle contractility measurements, muscle force values were normalized to each colonic ring’s mass. Sigmoidal dose-response curves were fitted with nonlinear regression, and best-fit parameters (logEC50, top bound, expressed as value ± SEM) between groups were compared using extra sum-of-squares F test in Prism 4.0 (Graphpad, San Diego, CA).

## Acknowledgments

We are grateful to V. Lennon (Mayo Clinic) for the ANNA-1 antisera, M. Shen (Columbia University) for AR^flx/flx^ mice, as well as V. Cheng (Beth Israel Deaconess Medical Center; BIDMC) and other members of the BIDMC GI Division for clinical trial support. We thank D. Ginty, D. Breault, M. Rutlin and J. Silvester for critical reading of the manuscript. This project was supported by funding from the American Gastroenterological Association-Takeda Research Scholar Award (M.R.), NIH K08DK125636 (M.R.), NIH R03DK110532 (M.R.), and NIH R01AT008573 (A.L.). Core facilities used for experiments were supported by the Harvard Digestive Disease Center (NIH P30DK034854) and the Boston Children’s Hospital/Harvard Medical School Intellectual and Developmental Disabilities Research Center (NIH U54 HD090255).

## Author contributions

D.R. and M.R. conceived the study and wrote the initial manuscript. D.R., A.R., L.T.M, K.P., A.P., M.M., W. D., P.Y. and M.R. performed experiments and analyzed results. R.H., S.B., K.H. and M.R. performed the analysis of the human sex hormone data. M.R., K.H., J.N. and A.L obtained funding for the studies. All authors participated in revising and approving the final manuscript. The authors declare no competing interests.

## Reference

1. Oliva, M., et al. The impact of sex on gene expression across human tissues. Science (New York, N.Y.) 369(2020).

2. Goodman, W.A., Erkkila, I.P. & Pizarro, T.T. Sex matters: impact on pathogenesis, presentation and treatment of inflammatory bowel disease. Nature reviews. Gastroenterology & hepatology (2020).

3. Camilleri, M. Sex as a biological variable in irritable bowel syndrome. Neurogastroenterol Motil 32, e13802 (2020).

4. Meoni, S., Macerollo, A. & Moro, E. Sex differences in movement disorders. Nature reviews. Neurology 16, 84–96 (2020).

5. Looijer-van Langen, M., et al. Estrogen receptor-β signaling modulates epithelial barrier function. Am J Physiol Gastrointest Liver Physiol 300, G621–626 (2011).

6. Jiang, Y., Greenwood-Van Meerveld, B., Johnson, A.C. & Travagli, R.A. Role of estrogen and stress on the brain-gut axis. Am J Physiol Gastrointest Liver Physiol 317, G203–G209 (2019).

7. D'Errico, F., et al. Estrogen receptor β controls proliferation of enteric glia and differentiation of neurons in the myenteric plexus after damage. Proc Natl Acad Sci U S A 115, 5798–5803 (2018).

8. Foradori, C.D., Weiser, M.J. & Handa, R.J. Non-genomic actions of androgens. Frontiers in neuroendocrinology 29, 169–181 (2008).

9. Winborn, W.B., Sheridan, P.J. & McGill, H.C., Jr. Sex steroid receptors in the stomach, liver, pancreas, and gastrointestinal tract of the baboon. Gastroenterology 92, 23–32 (1987).

10. Gonzalez-Montelongo, M.C., Marin, R., Gomez, T. & Diaz, M. Androgens differentially potentiate mouse intestinal smooth muscle by nongenomic activation of polyamine synthesis and Rho kinase activation. Endocrinology 147, 5715–5729 (2006).

11. Gonzalez-Montelongo, M.C., Marin, R., Gomez, T., Marrero-Alonso, J. & Diaz, M. Androgens induce nongenomic stimulation of colonic contractile activity through induction of calcium sensitization and phosphorylation of LC20 and CPI-17. Molecular endocrinology 24, 1007–1023 (2010).

12. Ortiz, H.E. & Cavicchia, J.C. Androgen-induced changes in nuclear pore number and in tight junctions in rat seminal vesicle epithelium. The Anatomical record 226, 129–134 (1990).

13. Furness, J.B., Callaghan, B.P., Rivera, L.R. & Cho, H.J. The enteric nervous system and gastrointestinal innervation: integrated local and central control. Advances in experimental medicine and biology 817, 39–71 (2014).

14. Corsetti, M., et al. First translational consensus on terminology and definitions of colonic motility in animals and humans studied by manometric and other techniques. Nature reviews. Gastroenterology & hepatology 16, 559–579 (2019).

15. Swaminathan, M., et al. Video Imaging and Spatiotemporal Maps to Analyze Gastrointestinal Motility in Mice. Journal of visualized experiments: JoVE, 53828 (2016).

16. Sanders, K.M., Koh, S.D., Ro, S. & Ward, S.M. Regulation of gastrointestinal motility--insights from smooth muscle biology. Nature reviews. Gastroenterology & hepatology 9, 633–645 (2012).

17. McClain, J.L., et al. Ca2+ responses in enteric glia are mediated by connexin-43 hemichannels and modulate colonic transit in mice. Gastroenterology 146, 497–507 e491 (2014).

18. Rao, M., et al. Enteric Glia Regulate Gastrointestinal Motility but Are Not Required for Maintenance of the Epithelium in Mice. Gastroenterology 153, 1068–1081 e1067 (2017).

19. Hao, M.M., et al. Early development of electrical excitability in the mouse enteric nervous system. J Neurosci 32, 10949–10960 (2012).

20. Mauvais-Jarvis, F. Sex differences in metabolic homeostasis, diabetes, and obesity. Biology of sex differences 6, 14 (2015).

21. Juntti, S.A., et al. The androgen receptor governs the execution, but not programming, of male sexual and territorial behaviors. Neuron 66, 260–272 (2010).

22. Jean-Faucher, C., Berger, M., de Turckheim, M., Veyssiere, G. & Jean, C. Developmental patterns of plasma and testicular testosterone in mice from birth to adulthood. Acta endocrinologica 89, 780–788 (1978).

23. Lewis, A.E., Vasudevan, H.N., O'Neill, A.K., Soriano, P. & Bush, J.O. The widely used Wnt1-Cre transgene causes developmental phenotypes by ectopic activation of Wnt signaling. Developmental biology 379, 229–234 (2013).

24. Drokhlyansky, E., et al. The Human and Mouse Enteric Nervous System at Single-Cell Resolution. Cell 182, 1606–1622 e1623 (2020).

25. Houghton, L.A., Jackson, N.A., Whorwell, P.J. & Morris, J. Do male sex hormones protect from irritable bowel syndrome? The American journal of gastroenterology 95, 2296–2300 (2000).

26. Kim, B.J., et al. Male sex hormones may influence the symptoms of irritable bowel syndrome in young men. Digestion 78, 88–92 (2008).

27. Ballou, S., et al. Open-label versus double-blind placebo treatment in irritable bowel syndrome: study protocol for a randomized controlled trial. Trials 18, 234 (2017).

28. Penson, D.F., Ng, C., Cai, L., Rajfer, J. & González-Cadavid, N.F. Androgen and pituitary control of penile nitric oxide synthase and erectile function in the rat. Biology of reproduction 55, 567–574 (1996).

29. Markle, J.G., et al. Sex differences in the gut microbiome drive hormone-dependent regulation of autoimmunity. Science (New York, N.Y.) 339, 1084–1088 (2013).

30. Colldén, H., et al. The gut microbiota is a major regulator of androgen metabolism in intestinal contents. Am J Physiol Endocrinol Metab 317, E1182–E1192 (2019).

31. Dart, D.A., Waxman, J., Aboagye, E.O. & Bevan, C.L. Visualising androgen receptor activity in male and female mice. PloS one 8, e71694 (2013).

32. Harman, S.M., Metter, E.J., Tobin, J.D., Pearson, J. & Blackman, M.R. Longitudinal effects of aging on serum total and free testosterone levels in healthy men. Baltimore Longitudinal Study of Aging. J Clin Endocrinol Metab 86, 724–731 (2001).

## Methods References

1. Rao, M., et al. Enteric glia express proteolipid protein 1 and are a transcriptionally unique population of glia in the mammalian nervous system. Glia 63, 2040–2057 (2015).

2. Rao, M., et al. Enteric Glia Regulate Gastrointestinal Motility but Are Not Required for Maintenance of the Epithelium in Mice. Gastroenterology 153, 1068–1081 e1067 (2017).

3. Swaminathan, M., et al. Video Imaging and Spatiotemporal Maps to Analyze Gastrointestinal Motility in Mice. Journal of visualized experiments: JoVE, 53828 (2016).

4. Francis, C.Y., Morris, J. & Whorwell, P.J. The irritable bowel severity scoring system: a simple method of monitoring irritable bowel syndrome and its progress. Alimentary pharmacology & therapeutics 11, 395–402 (1997).

