## Extended Data Figures 1-8 for "Androgen regulation of bowel function in mice and humans"

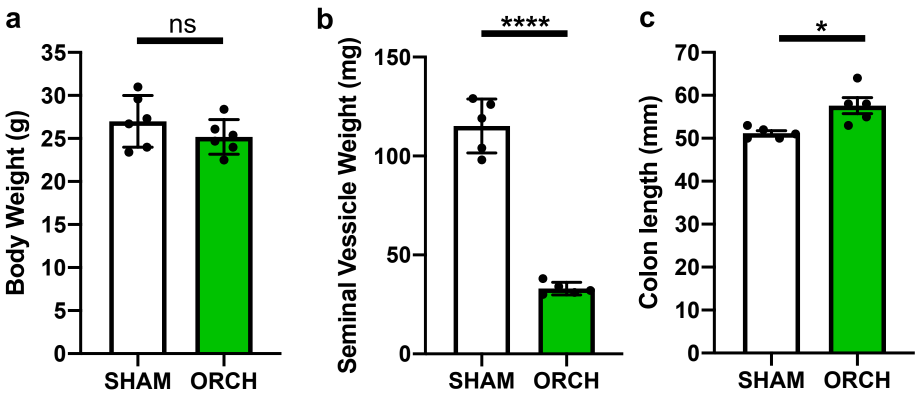


**Extended Data Figure 1. Loss of gonadal function alters androgen-responsive tissues but not body weight.**
**a.** Body mass measured in 10 week old mice, four weeks after bilateral orchiectomy (ORCH) or sham surgery (SHAM), was no different between the groups (*N* = 6 mice/group).
**b.** Mass of seminal vesicles, a tissue well-known to atrophy in the absence of androgens, is markedly lower in ORCH than SHAM mice 4 weeks after surgery (*N* = 5 mice/group).
**c**. Colon lengths in ORCH mice are 10% greater compared to SHAM mice, 4-6 weeks after surgery (*N* = 5 mice/group).
* Represents $P < 0.05$, ** $P < 0.01$, *** $P < 0.005$, and **** $P < 0.001$.

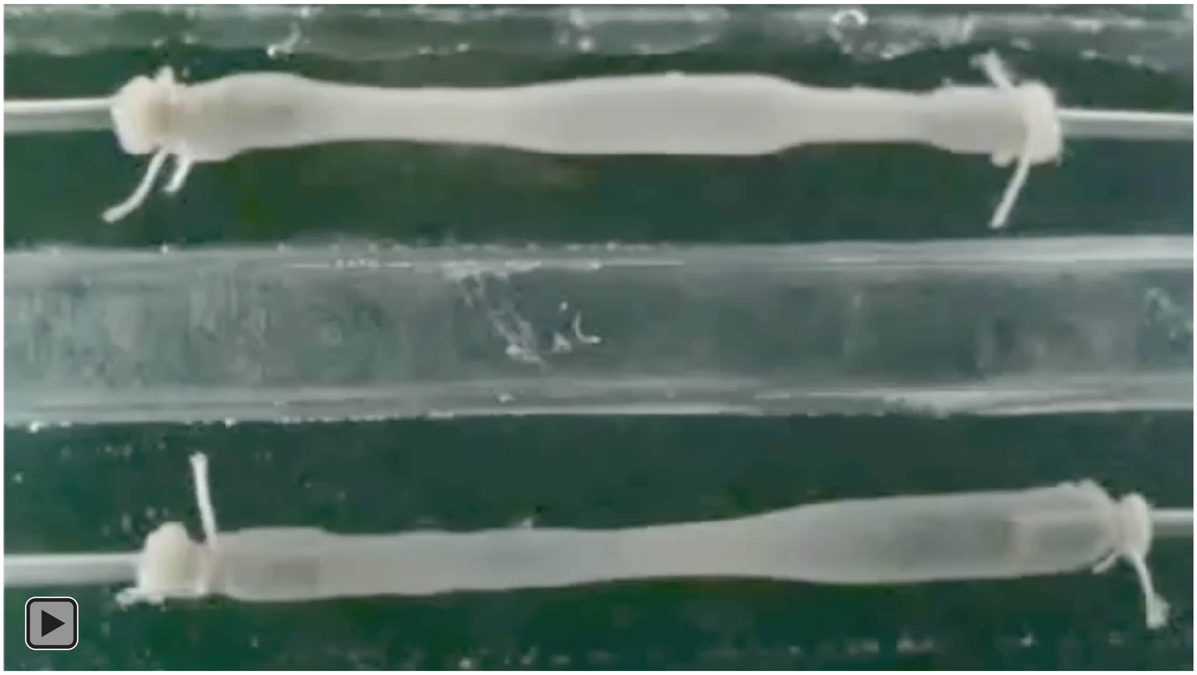


**Extended Data Figure 2. Colonic contractile activity is disorganized and less effective at oral-to-anal propulsion of luminal contents in male mice lacking gonadal function.**

**Video-recordings of motor activity in colons from 10 week old mice four weeks after sham operation (SHAM, top bath) and bilateral orchiectomy (ORCH, bottom bath). Colons are oriented with oral end on the left and anal end on the right. The colon from the SHAM mouse shows distinct colonic migrating motor contractions (CMMCs) that progress from oral to anal ends of the colon, as typically seen in colons from healthy, adult mice. The colon from the ORCH mouse, in contrast, has numerous disorganized, irregular contractions that do not consistently progress from oral to anal ends. These recordings are representative of observations made in three 15 minute videos obtained from each of at least 5 mice per condition.**

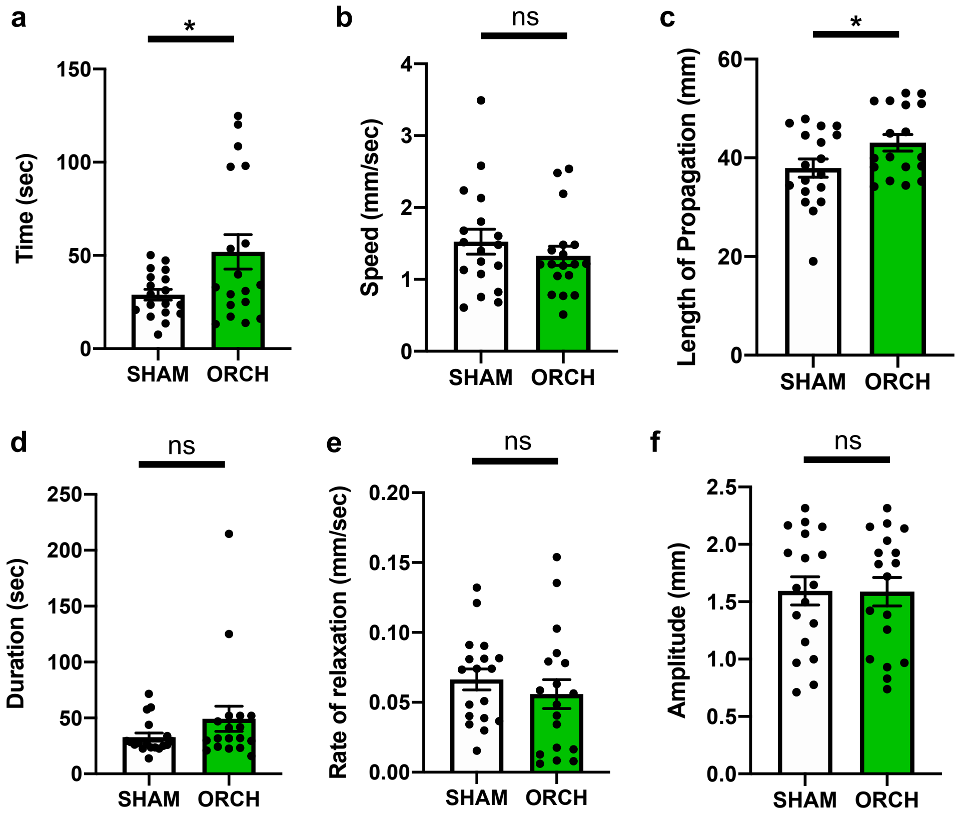


**Extended Data Figure 3. Gonadal androgen deficiency disrupts colonic motility.**
Features of colonic migrating motor contractions (CMMCs) in colons from SHAM and ORCH mice that were acutely isolated and imaged *ex vivo*, four or more weeks after surgery. CMMCs were defined as contractions that originated in the proximal colon and successfully propagated at least 50% of the length of the colon. All parameters were measured from three 15 minute video recordings obtained from each of 6 mice per group. Each individual data point is shown. The organ bath preparation accommodates 2 colons at a time, and SHAM and ORCH colons were imaged in every session in parallel. *P*-values reflect unpaired t-tests.

**a.** Time in seconds (sec) required for colons to relax from maximal contraction to baseline gut width following a CMMC was greater in colons from ORCH mice ($P = 0.034$).

**b.** CMMC speed was no different between colons from SHAM and ORCH mice ($P = 0.3755$).

**c.** Length of propagation of individual CMMCs was greater in colons from ORCH mice ($P = 0.0458$), compared to SHAM controls, proportional to the longer colonic length in ORCH mice (see Extended Data Fig. 1c).

**d.** Duration of CMMCs was no different between colons from SHAM and ORCH mice ($P = 0.1740$).

**e.** Rate of colonic relaxation following a CMMC was no different in colons from SHAM and ORCH mice ($P = 0.4152$).

**f.** Amplitude of gut contraction during CMMCs was no different in colons from SHAM and ORCH mice ($P = 0.9717$).

* Denotes $P < 0.05$. ns denotes $P > 0.05$.

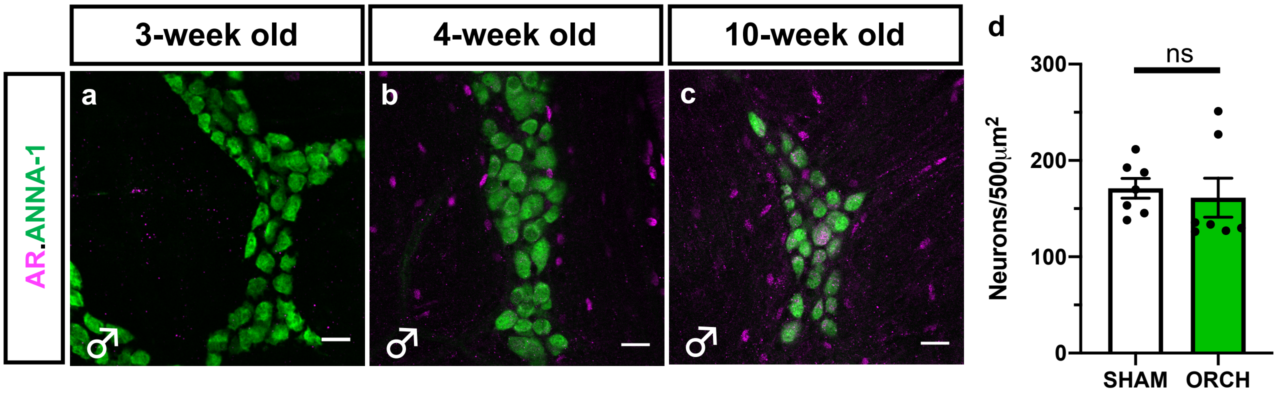


**Extended Data Figure 4. Androgen receptor is upregulated in enteric neurons following puberty and subsequent loss of male gonadal function does not alter enteric neuronal density.**

**a-c.** Androgen receptor (AR) immunoreactivity is undetectable in the muscularis externa of a colon from a 3-week old wildtype male mouse (**a**). By 4-weeks of age, AR is expression is detectable in scattered cells outside the myenteric plexus, but not in the ENS (**b**). By the post-pubertal age of 10 weeks, robust AR expression is observed both within a subset of enteric neurons and in the smooth muscle surrounding the myenteric plexus (**c**, see Main Figure 2g for an additional example.) Enteric neuronal soma are marked by ANNA-1 immunoreactivity. Scale bar = 50µm.

**b.** Neuronal density in the myenteric plexus, measured by whole mount immunostaining of colonic segments from SHAM and ORCH mice for ANNA-1, shows that loss of gonadal function does not alter enteric neuronal number five weeks after surgery ($N$ = 7 mice/group).

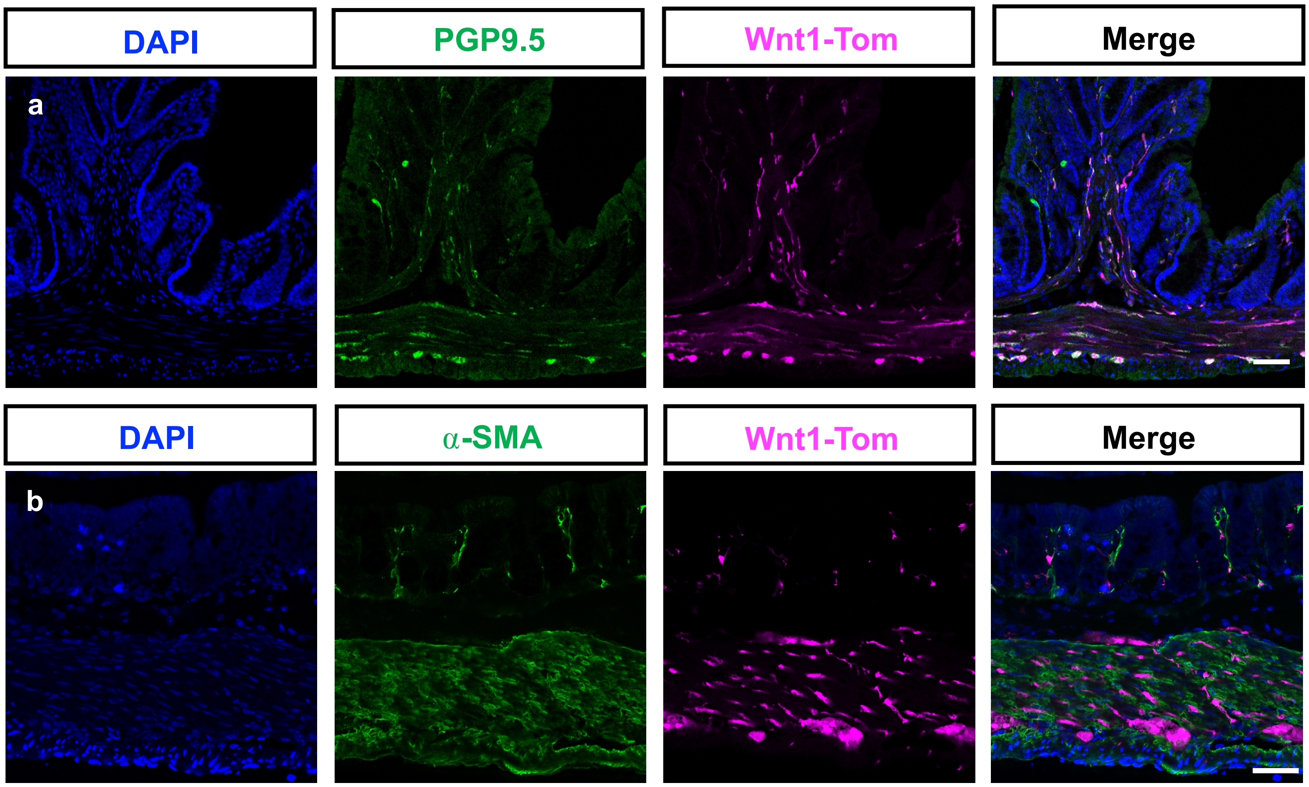


**Extended Data Figure 5. Wnt1-Cre2 transgenic mice exhibit Cre activity in enteric neurons, but not smooth muscle, in the adult mouse colon.**

$Wnt1^{Cre2}$::$Rosa26^{Ai9/+}$ mice were analyzed to assess Cre activity in the colon. Cre activity induces expression of a tdTomato fluorescent reporter protein in Ai9 mice. Representative cross-sections of colons obtained from 10-week old mice are shown. Cross-sections immunostained for the pan-neuronal marker PGP9.5 (panels in row **a**) show that PGP9.5 colocalizes with the tdTomato reporter, confirming Cre activity in the majority of enteric neurons. In contrast, the smooth muscle marker α-smooth muscle actin (SMA) does not colocalize with tdTomato (panels in row **b**), indicating that there is no Cre-mediated recombination in the colonic smooth muscle. Nuclei counterstained with DAPI. Scale bars = 50µm

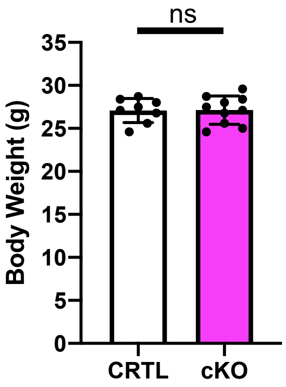


**Extended Data Figure 6. Genetic depletion of androgen signaling in the peripheral nervous system does not alter body weight.**

Body mass measured in 10 week old $AR^{Wnt1KO}$ mice (cKO) and littermate controls (CTRL; $Ar^{flx/Y}$) was no different between the groups ($N$ = 8-10 mice/group). "ns" represents $P$ = 0.9545 resulting from unpaired t-test comparing two group means.

| | MEAN AGE | SD | MEAN IBS-SSS | SD |
|---|---|---|---|---|
| Male | | | | |
| IBS-All Subtypes (58) | 43.19 | 17.29 | 280.36 | 66.30 |
| IBS-C (14) | 50.43 | 22.84 | 273.07 | 65.29 |
| IBS-D (41) | 41.93 | 14.69 | 278.49 | 67.23 |
| IBS-M (1) | 27.00 | . | 374.00 | . |
| IBS-U (2) | 26.50 | 2.12 | 323.00 | 32.53 |
| Healthy Controls (14) | 34.79 | 10.65 | - | - |
| Female | | | | |
| IBS-All Subtypes (150) | 43.34 | 19.16 | 271.28 | 63.69 |
| IBS-C (59) | 40.64 | 18.14 | 285.12 | 65.45 |
| IBS-D (66) | 46.77 | 20.50 | 258.62 | 60.73 |
| IBS-M (22) | 42.23 | 17.32 | 276.09 | 59.73 |
| IBS-U (3) | 29.00 | 8.72 | 242.33 | 90.18 |
| Healthy Controls (14) | 38.64 | 13.74 | - | - |

**Extended Data Figure 7. Characteristics of human study participants.**
Mean age, IBS symptom severity score (IBS-SSS) and associated standard deviations (SD) for healthy control study participants, and participants with IBS categorized into individual IBS subtypes. IBS-Constipation (IBS-C), IBS-Diarrhea (IBS-D), IBS-Mixed presentation (IBS-M), and IBS-Undetermined (IBS-U).

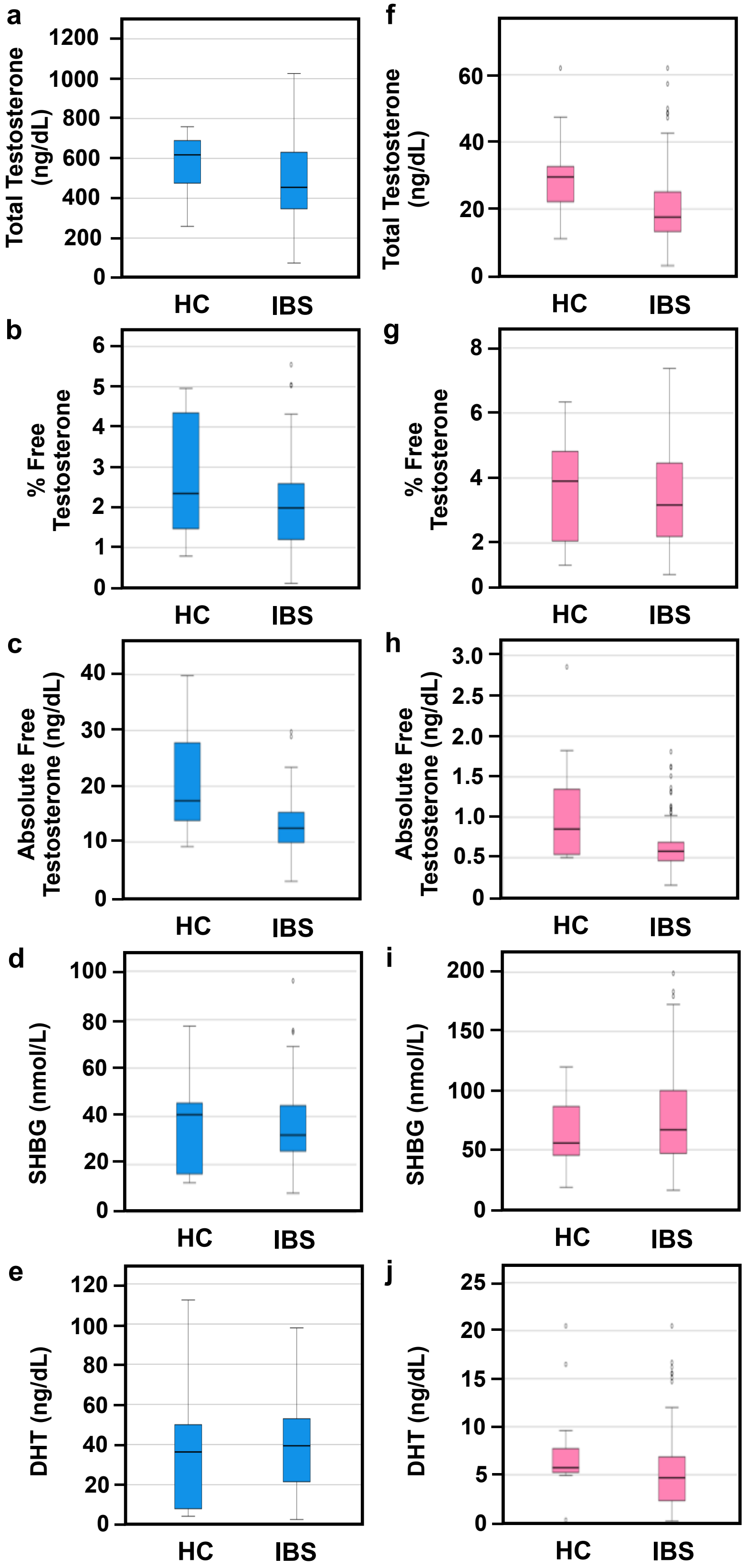


**Extended Data Figure 8. Distribution of androgen levels among IBS patients and healthy controls.**

Box-plots illustrating levels of total testosterone, percent free testosterone, absolute free testosterone, sex hormone binding globulin (SHBG) and DHT in healthy controls (HC) and patients with IBS.

**a – e**. Males

**f – j**. Females
